# Neurons in primate entorhinal cortex represent gaze position in multiple spatial reference frames

**DOI:** 10.1101/182220

**Authors:** Miriam L. R. Meister, Elizabeth A. Buffalo

**Affiliations:** Washington National Primate Research Center, Seattle, WA, 98195, United States; Department of Physiology and Biophysics, University of Washington, Seattle, WA, 98195, United States; University of Washington School of Medicine, Seattle, WA,98195, United States

## Abstract

Primates predominantly rely on vision to gather information from the environment, and neurons representing visual space and gaze position are found in many brain areas. Within the medial temporal lobe, a brain region critical for memory, neurons in the entorhinal cortex of macaque monkeys exhibit spatial selectivity for gaze position. Specifically, the firing rate of single neurons reflects fixation location within a visual image (Killian et al., 2012). In the rodents, entorhinal cells such as grid cells, border cells, and head direction cells show spatial representations aligned to visual environmental features instead of the body (Hafting et al., 2005, Solstad et al. 2008, Sargolini et al., 2006, Diehl et al., 2017). However, it is not known whether similar allocentric representations exist in primate entorhinal cortex. Here, we recorded neural activity in the entorhinal cortex in two male rhesus monkeys during a naturalistic, free-viewing task. Our data reveal that a majority of entorhinal neurons represent gaze position, and that simultaneously recorded neurons exhibit distinct spatial reference frames, with some neurons aligning to the visual image and others aligning to the monkey’s head position. Our results also show that entorhinal neural activity can be used to predict gaze position with a high degree of accuracy. These findings demonstrate that visuospatial representation is a fundamental property of entorhinal neurons in primates, and suggest that entorhinal cortex may support relational memory and motor planning by coding attentional locus in distinct, behaviorally relevant frames of reference.

**Significance Statement:** The entorhinal cortex, a brain area important for memory, shows striking spatial activity in rodents through grid cells, border cells, head direction cells, and nongrid spatial cells. The majority of entorhinal neurons signal the location of a rodent relative to visual environmental cues, representing the location of the animal relative to space in the world instead of the body. Recently, our laboratory found that entorhinal neurons can signal location of gaze while a monkey visually explores images. Here, we report that spatial entorhinal neurons are widespread in the monkey, and these neurons are capable of showing a world-based spatial reference frame locked to the bounds of explored images. These results help connect the extensive findings in rodents to the primate.

## Introduction

The rodent entorhinal cortex contains cell types that represent body position, including grid cells, border cells, and head direction cells (Hartley et al., 2013). However, unlike many sensory and motor areas of the brain that represent spatial information relative to the body (egocentric representation), entorhinal cells fire when the body occupies a particular space relative to features in the local environment (allocentric representation) (Quirk et al., 1992; Fyhn et al., 2004; Hafting et al., 2005; Sargolini et al., 2006; Barry et al., 2007; Fyhn et al., 2007; Savelli et al., 2008; Aronov and Tank, 2014). For example, an entorhinal neuron will fire selectively when a rodent is located in a particular place within an enclosure that has a visible cue card on the wall, and then fire for that same location relative to the card when the card is moved (Quirk et al., 1992). This world-based, allocentric reference frame contrasts with the egocentric spatial reference frame of sensory and motor neurons. Specifically, while visual cortical neurons fire egocentrically to reflect the spatial arrangement of stimuli on the retina, entorhinal neurons instead fire allocentrically across highly variable visual inputs (Quirk et al., 1992), or even without visual input (Hafting et al., 2005), in order to reflect the same body position relative to objects in the world.

The great majority of this rich line of work on entorhinal spatial coding has been conducted in rodents, leaving open the question of whether primate entorhinal neurons display allocentric coding, and how such coding might support the memory function of this brain area. The neuronal algorithms that enable episodic memory are likely to be exposed using these spatial representations as a window into circuit computations. For example, study of these spatial representations may reveal the neural basis of spatially mnemonic behavior, such as using an allocentric coordinate frame to direct gaze to the location next to the refrigerator, where you recall leaving your keys. Allocentric representations have been identified in other regions of the human and nonhuman primate brain, e.g., (Olson and Gettner, 1995; Georges-François et al., 1999; Committeri et al., 2004; Chen and Crawford, 2017). Recent work demonstrated that primate entorhinal cells exhibit spatial representations by firing selectively when a monkey looked at certain regions of space on a computer screen while freely viewing images (Killian et al., 2012). However, because the image location was held constant in those experiments, it was unclear whether primate entorhinal neurons reflected gaze position relative to the skull in egocentric space, or instead reflected allocentric space locked to conspicuous environmental features, such as the image boundaries. In order to address this question, and to assess the proportion of neurons with spatial representations, we recorded gaze position and the activity of individual entorhinal neurons in monkeys freely viewing images that were displayed in different screen locations across trial blocks. We determined if neurons represented where the monkey was looking, and importantly, whether gaze position was coded relative to the visual image position or relative to the monkey’s head. The results identified a strikingly large proportion of entorhinal cells that coded gaze position, and while some cells coded allocentric space by showing consistent spatial firing locked to the image display window across its varied screen locations, other simultaneously recorded cells did not alter their spatial firing when the location of the image display window was changed. Taken together, these data show that gaze position is represented widely across the primate entorhinal cortex, and gaze position is represented in multiple spatial reference frames simultaneously across the neuronal population.

## Materials and Methods

### Experimental design and statistical analyses

In order to identify the frame of reference for spatial entorhinal neurons, we recorded gaze position and the spiking activity from single entorhinal neurons while head-stabilized rhesus macaque monkeys freely viewed large, complex images that were presented at two different locations on a stationary screen within the recording session (Figure 1). In Left trial blocks, images were centered 2° to the left of the center of the screen, whereas in Right trial blocks, images were centered 2° to the right of the center of the screen, resulting in a total offset of 4° between the image locations. Data were obtained from two rhesus macaque monkeys.

**Figure 1.**
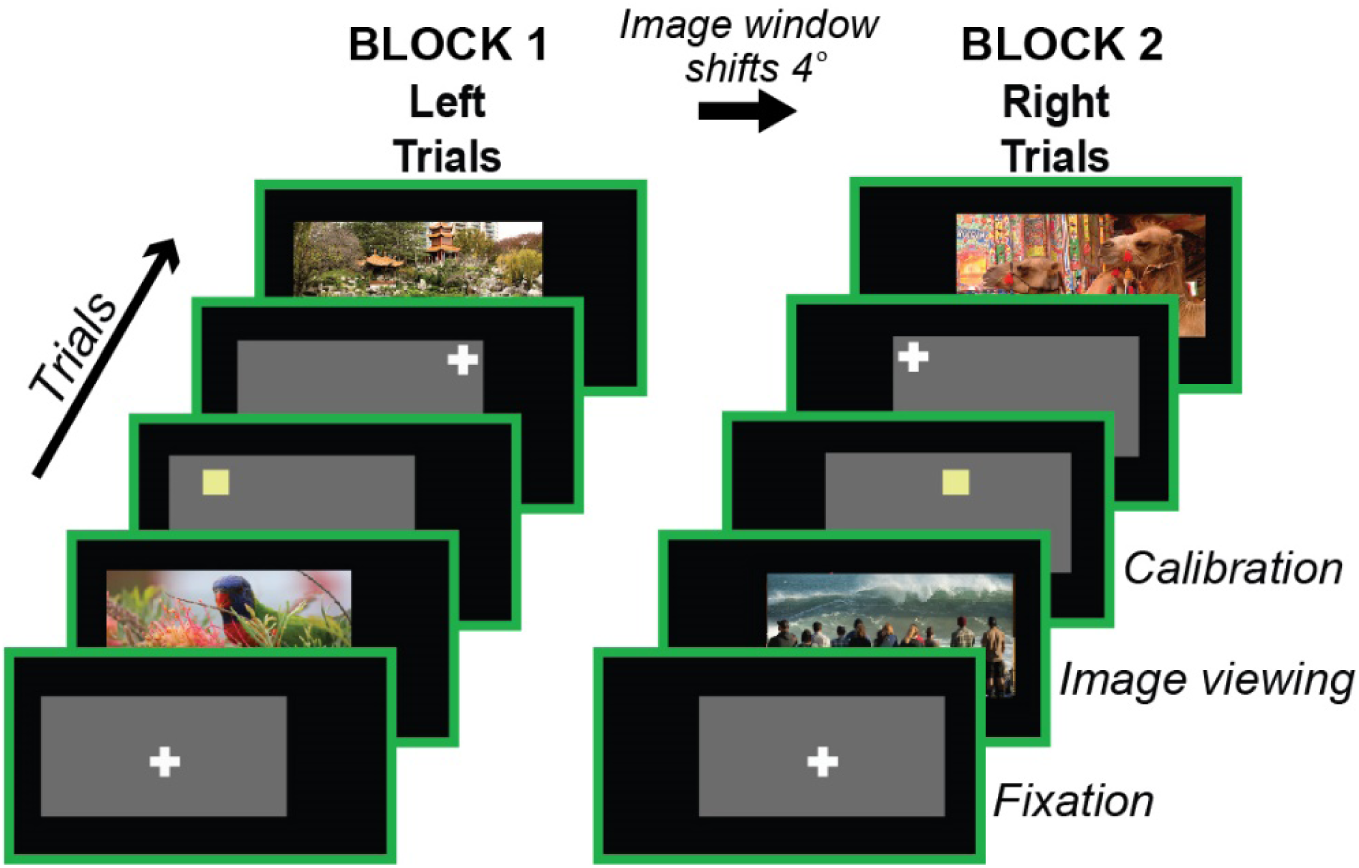
Free-viewing paradigm. *Task schematic.* Left and Right trial blocks differed by a 4° shift in visual stimulus location relative to the screen borders (green rectangle). In Left trials, stimuli were centered 2° left of the screen center, whereas in Right trials, stimuli were centered 2° to the right of the screen center. The monkey’s head remained in the same position relative to the room and the computer screen for all trials. All stimuli presented within each trial block were confined within the screen space occupied by the images of that block (the “image window” of that trial block). *Fixation:* Trials began with a 500 - 750 ms required fixation on a cross positioned pseudorandomly in one of 9 potential locations across a gray background rectangle. *Image Viewing:* A complex image (photograph of variable content) was displayed for 5 seconds of free viewing. *Calibration:* The monkey received a fruit slurry reward for releasing a response bar in response to a subtle color change of small square (which could appear in multiple locations across the gray background rectangle). The monkey’s gaze on this square was used to calibrate eye tracking software, and correct any drift in recorded eye position. A minimum of 30 image trials were presented within each block. Right and Left trial blocks started alternate recording sessions.

Non-parametric bootstrapping procedures were used to determine significance at the level of p ≤ 0.05 for all statistical analyses unless otherwise noted. Details of how bootstrapping was used in different analyses are reported along with the description of each analysis. Unless specified, all analyses were performed using custom code in MATLAB (The MathWorks, Natick, MA).

### Subjects, training and surgery

Two male rhesus macaques, 10 and 11 years old, and weighing 13.8 and 16.7 kg respectively, were trained to sit in a primate chair (Crist Instrument Company, Inc., Hagerstown, MD) with a fixed head position and to release a touch-bar for fruit slurry reward delivered through a tube. The monkeys were trained to perform various tasks by releasing the touch-bar at appropriate times relative to visual stimuli presented on a screen. Magnetic resonant images of each monkey’s head were made both before and after surgery to plan and confirm implant placement. Separate surgeries were performed to implant a head post, then months later, a recording chamber, and finally a craniotomy within the chamber. All experiments were performed in accordance with protocols approved by the Emory University and University of Washington Institutional Animal Care and Use Committees.

### Behavioral task

For all recordings, the monkey was seated in a dark room, head fixed and positioned so that the center of the screen (54.1 cm × 29.9 cm LCD screen, 120 Hz refresh rate, 1280 × 720 pixels, BenQ America Corp., Costa Mesa, CA) was aligned with his neutral gaze position and 60 cm away from the plane of the his eyes (equating to ∼25 screen pixels per degree of visual angle, or 1°/ cm). Stimulus presentation was controlled by a PC running Cortex software (National Institute of Mental Health, Bethedsa, MD). Gaze location was monitored at 240 Hz with an infrared eye-tracking system (I-SCAN, Inc., Woburn, MA).

Gaze location was calibrated before and during each recording session with calibration trials (Figure 1), in which the monkey held a touch-sensitive bar while fixating a small (0.5°) gray square presented at various locations on the monitor. The square turned yellow 400 - 750 ms (uniform distribution) after its appearance, and the monkey was required to release the bar in response to the color change for delivery of the fruit slurry reward. The subtlety of the color change forced the monkey to visually fixate the location of the small square to correctly perform those trials, therefore allowing calibration of gaze position to the displayed stimuli. Specifically, the gain and offset of the recorded gaze position were adjusted so that gaze position matched the position of the fixated stimulus. Throughout the session, calibration trials enabled continual monitoring of the quality of gaze position data and correction of any drift. The monkeys performed alternating blocks of trials in which presented images were either centered slightly to the right or centered slightly to the left (Figure 1). The left and right image window locations were offset by 4°. Before each image presentation, a crosshair (0.3° × 0.3°) appeared in one of nine possible locations across a gray background rectangle superimposed on the dark background of the screen. The gray rectangle was the same size and position of the images presented in that trial block, and encompassed the screen space for all visual stimuli in that trial block. Once gaze position registered within a 3° × 3° window around the crosshair, and was maintained within that spatial window for 500 - 750 ms, then the image was presented. Images were complex natural images downloaded from the public photo-sharing website, Flickr (www.flickr.com). If necessary, images were re-sized by the experimenter for stimulus presentation (sized 30° × 15° for Monkey WR and 30° × 25° for Monkey MP). Monkeys were allowed to freely view the image, and the image vanished after gaze position had registered within the image frame for a cumulative 5 seconds. No food reward was given during image viewing trials. Each image presentation was followed by three calibration trials. The color-change square of calibration trials was superimposed on the gray background rectangle of a given image block. To adequately cover the relevant screen space for calibration during and after the experiment, there were a large number of unique color-change square locations (100 and 54 unique locations for Monkey MP and Monkey WR, respectively) across image-location trial blocks, as well as an additional 18 unique calibration points from the required fixation on the crosshair. These frequent and spatially-ranging calibration points ensured that gaze position data could be tracked for accuracy across the entire session for the whole expanse of relevant screen area.

After completing a block of trials, a new block of trials would begin with all visual stimuli (images and calibration trial stimuli) shifted laterally 4°. Stimuli in the Left trial block were centered 2° to the left of the center of the screen, whereas stimuli in the Right trial block were centered 2° to the right of the center of the screen. Left and Right trial blocks were pseudorandomly selected to be the first trial block of an experimental session.

In the first 15 sessions (of 26 sessions total) for Monkey WR, an ABA design with three trial blocks was used (visual stimuli were centered the same way in the first and last trial blocks, whereas stimuli in the middle block were centered at a shifted location), with a total maximum of 180 image presentations per session. In the rest of the sessions for Monkey WR and all 14 sessions for Monkey MP, there was a total maximum of 240 image presentations across only two trial blocks (120 image presentations within each image window location).

### Offline eye position calibration

Eye position data from calibration trials were examined offline after the experiment in order to further improve calibration, and ensure fixation locations were stable throughout the experiment. Eye position traces during calibration trials were fit to the calibration points (and crosshair fixation data to crosshair points), with either an affine, polynomial, projective, or linear transformations in Matlab (“cp2tform” function). The best fitting transform was selected by visual inspection of plots showing calibration points and the fit eye position data from calibration. The selected transform was then applied to the rest of the eye position data. One session was excluded from further analysis because this check revealed an unsalvageable compromise in quality of the eye position data.

### Electrophysiology

Each recording session, a laminar electrode array (AXIAL array with 13 channels, FHC, Inc.) mounted on a microdrive (FHC, Inc.) was slowly lowered into the brain through the craniotomy. Magnetic resonant images along with the neural signal were used to guide the penetration. Spikes and local field potentials were recorded using hardware and software from Blackrock, Inc., and neural data were sampled at 30 kHz. A 500 Hz high-pass filter was applied, as well as an electric line cancellation at 60 Hz. In some recording sessions, a channel without any spiking activity was used as a reference electrode in order to subtract artifact noise (e.g., reward delivery, movement of the monkey). Spikes were sorted offline into distinct clusters using principle components analysis (Offline Sorter, Plexon, Inc.). Sorted clusters were then processed further by custom code in Matlab to eliminate any data where minimum inter-spike interval was less than 1 ms, and to identify any missed changes in signal (e.g., shrinking of the waveform of interest, a new waveform appearing), using a raster and plots of waveforms across the session for each cell. When change in signal was identified, appropriate cuts were made to exclude compromised spike data from before or after a change point. 455 potential single units originally cut in Offline Sorter were reduced to 357 single units. To further ensure recording location within the entorhinal cortex and identify from which cortical layers units were recorded, we examined each session’s data for the stereotypical, electrophysiological signature produced across entorhinal cortical layers at the onset of saccadic eye movement (Figure 1-1) (Killian et al., 2012; Killian et al., 2015). One recording session, which other electrode placement metrics suggest was conducted above the entorhinal cortex within the hippocampus, lacked this electrophysiological signature and was excluded from further analysis (8 single units were excluded from being categorized as entorhinal cells). No recording sessions showed the current source density electrophysiological signature of adjacent perirhinal cortex (Takeuchi et al., 2011) at stimulus onset. The laminar location of each recorded channel was estimated using approximate cortical thickness along with layer-specific signal features: The phase reversal across cortical layers that occurs near layer II ∼200 ms after the saccadic onset, and a phase reversal 100-150 ms after the saccadic onset indicating the transition to white matter dorsally. When one of these two, laminar-specific signals were missing, then the resulting ambiguities were retained in laminar classification (i.e., a neuron was classified as ambiguously being either in superficial or deep layers, as shown in the middle column of data in Figure 4-1A).

Location of cells along the anterior-posterior anatomical axis was accomplished by visually matching the anatomical features within a brain atlas (Paxinos et al., 2000) to the post-chamber implant surgery coronal MRI slice (1 mm slices) estimated to be the plane of a recording. The distance of cells from the rhinal sulcus was determined by the voxel distance (0.5 mm voxels) between the sulcus and the estimated recording location on a coronal MRI slice.

### Rate maps to characterize neural representation of gaze

In order identify neural activity related to gaze position as the monkey viewed the images, firing rate maps were computed for each neuron. The firing rate maps showed a neuron’s activity level across gaze positions within the screen space occupied by the image (Figure 2). Data from the first 500 ms of image viewing were excluded to avoid transient visual responses to the onset of the image. The image space was divided into 0.5° square spatial bins, and the number of spikes that occurred when the monkey’s eye position fell within each bin was divided by the total viewing time within that bin. In order to accommodate potentially different firing field sizes and levels of spatial resolution, rate maps were smoothed three different ways: Adaptive smoothing (Skaggs et al., 1993), and smoothing with a Gaussian filter (5.5° × 5.5°) that either had a standard deviation of 1° or 1.5°. All three of these smoothing methods produced similar-looking rate maps for a given piece of data. Although smoothing generated extrapolated values for unvisited spatial bins, these values were subsequently removed so that unvisited spatial bins were left empty.

**Figure 2.**
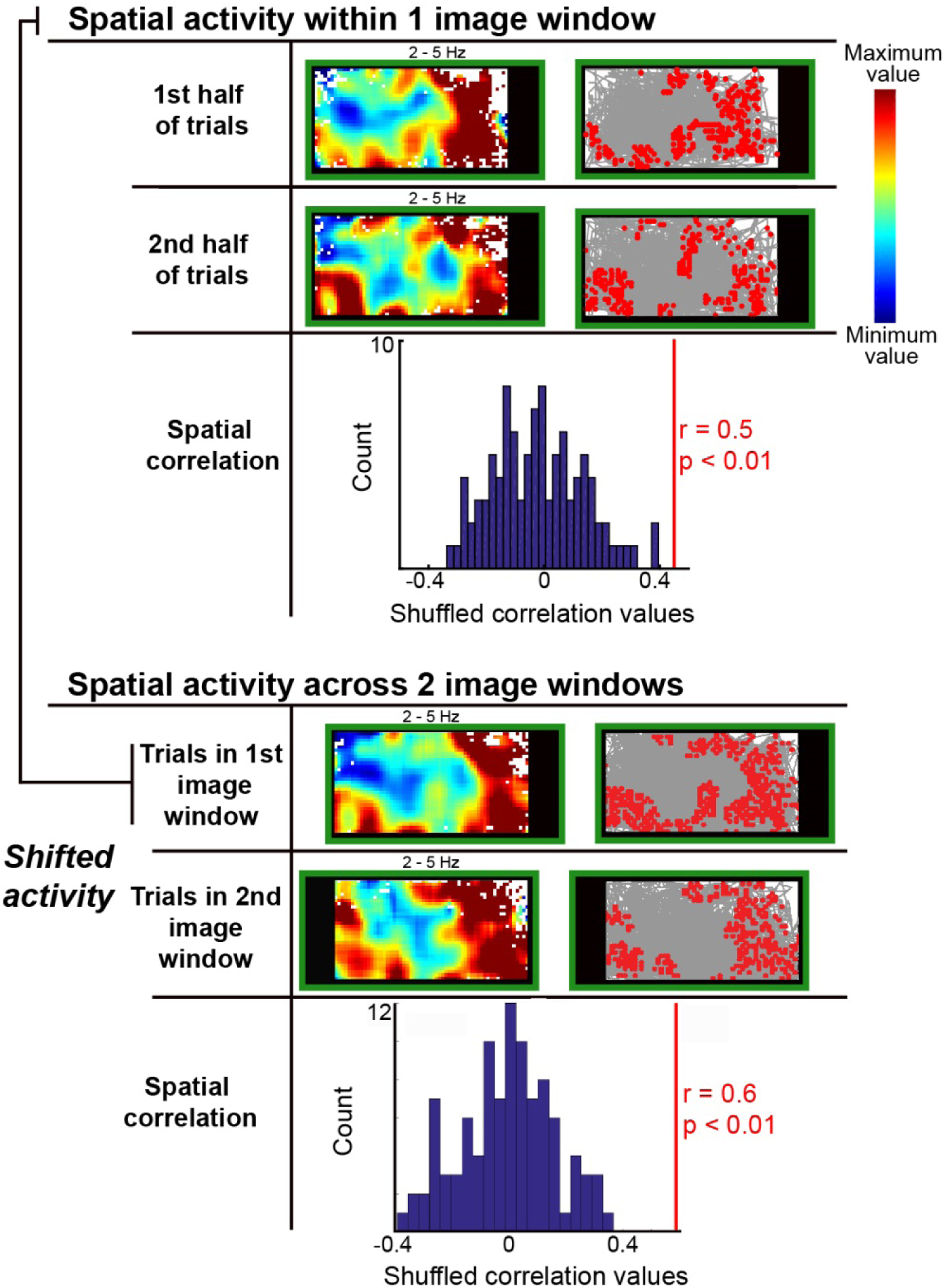
Spatial activity shifts with location of image window. Firing rate maps and the spatial correlation between them are shown for an example neuron with consistent spatial activity within one screen location (top) and across screen locations (bottom). The green rectangle schematically represents the borders of the screen, and the black represents screen space outside the image display window of a trial block. Warm and cool colors within the firing rate maps indicate high and low firing rate respectively; the firing rate reported on the top of each rate map corresponds to the minimum and maximum of the color bar (top right). To the right of each rate map, eye position trace is shown in gray, and red dots indicate gaze position locations where the neuron fired action potentials above the median firing rate. The histograms under each pair of rate maps show the spatial correlation value between the rate map pair (red line) relative to the shuffled correlation values (blue) that result when the spike train was shuffled relative to the eye position trace (100 shuffles for each rate map). Rate maps are the same size as presented images (30° × 15°). A movie of spikes occurring for this example cell as gaze position moves over the screen can be found in Extended Data (Movie 1).

### Spatial stability

To assess whether each neuron’s spatial activity was consistent for gaze position across trials, rate maps were assessed for stability across time. A spatial correlation (Pearson’s) was computed between two rate maps from one neuron from separate time periods of the neural recording. This spatial correlation was considered significant if it was higher than 95% of bootstrapped correlation values, which were computed from shuffled data of the neuron’s original two rate maps. Specifically, the spike train of a rate map was shifted circularly along the corresponding eye position trace at 100 equally spaced increments, i.e., the end of the spike train was wrapped to correspond to the beginning eye position data with the first shift. In order to avoid a large overlap with the original map, the starting positions of spike train shifts along the eye position trace were constrained to begin at least 10 seconds after the start of the trace and 10 seconds before the end of the trace. For each of the 100 shuffles, a new rate map was generated, and a spatial correlation was computed between the two generated rate maps. If the original spatial correlation was greater than or equal to the 95^th^ percentile of the 100 spatial correlation values generated from shuffling, then spatial correlation between the neuron’s two rate maps was considered to be significant (p ≤ 0.05), and the neuron was considered to show spatial stability (e.g., histogram in Figure 2). To determine spatial stability for a neuron across trials where images were presented in the *same* location, data from those trials were split across time to yield two rate maps that were then tested for correlated activity through the shuffling procedure described above (top panel of Figure 2). We then extended this analysis to examine spatial stability across trial blocks, when the image window was laterally shifted to a *different* location. In order to quantify spatial activity that shifted with the location of presented images, a correlation was computed for each neuron between the two rate maps from image viewing in the two different image display windows. A cell was considered to have a spatial representation that shifted along with the image location (“Shifted” spatial representation) if rate maps from the two different image window locations, aligned with the image bounds, yielded a significant spatial correlation (Figure 2). Likewise, a cell was considered to have a stable spatial representation that did *not* shift along with image window location (“Non-shifted” spatial representation) if rate maps from two different image window locations, but the same screen space, yielded a significant spatial correlation (Figure 3). The number of neurons falling into these two categories were assessed for each recording session (Figure 4).

**Figure 3.**
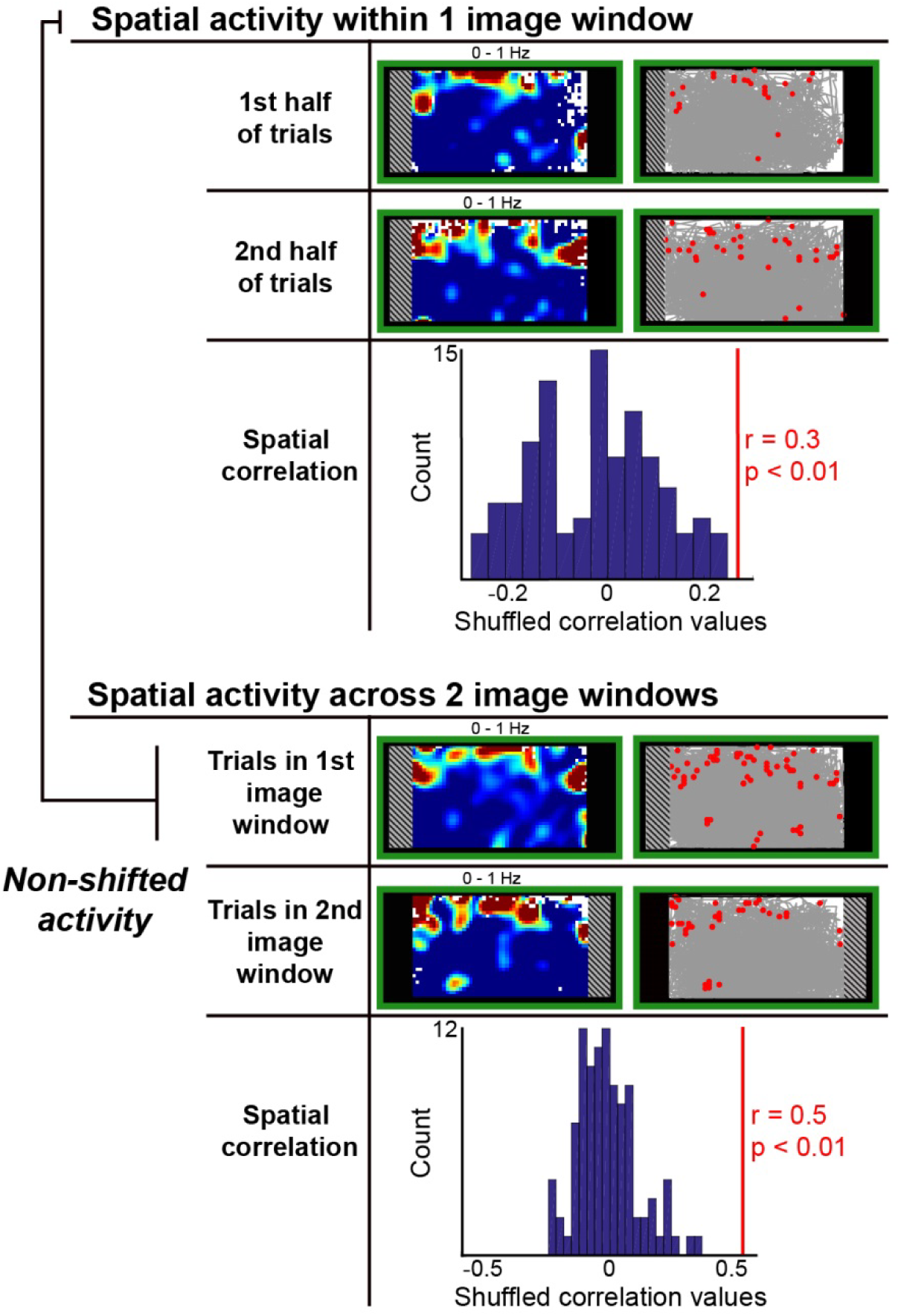
Example neuron with consistent spatial activity that does not shift with image window location. Firing rate maps and the spatial correlation between them are shown for a neuron that did not shift its activity with the location of image window. Plotting schematic is identical to Figure 2. Rate maps are the size of the image space common to all image presentations (26° × 15°), and crosshatching indicates image space of a trial block that was not covered by all images across all experimental trial blocks. A movie of spikes occurring for this example cell as gaze position moves over the screen can be found in Extended Data (Movie 2).

**Figure 4.**
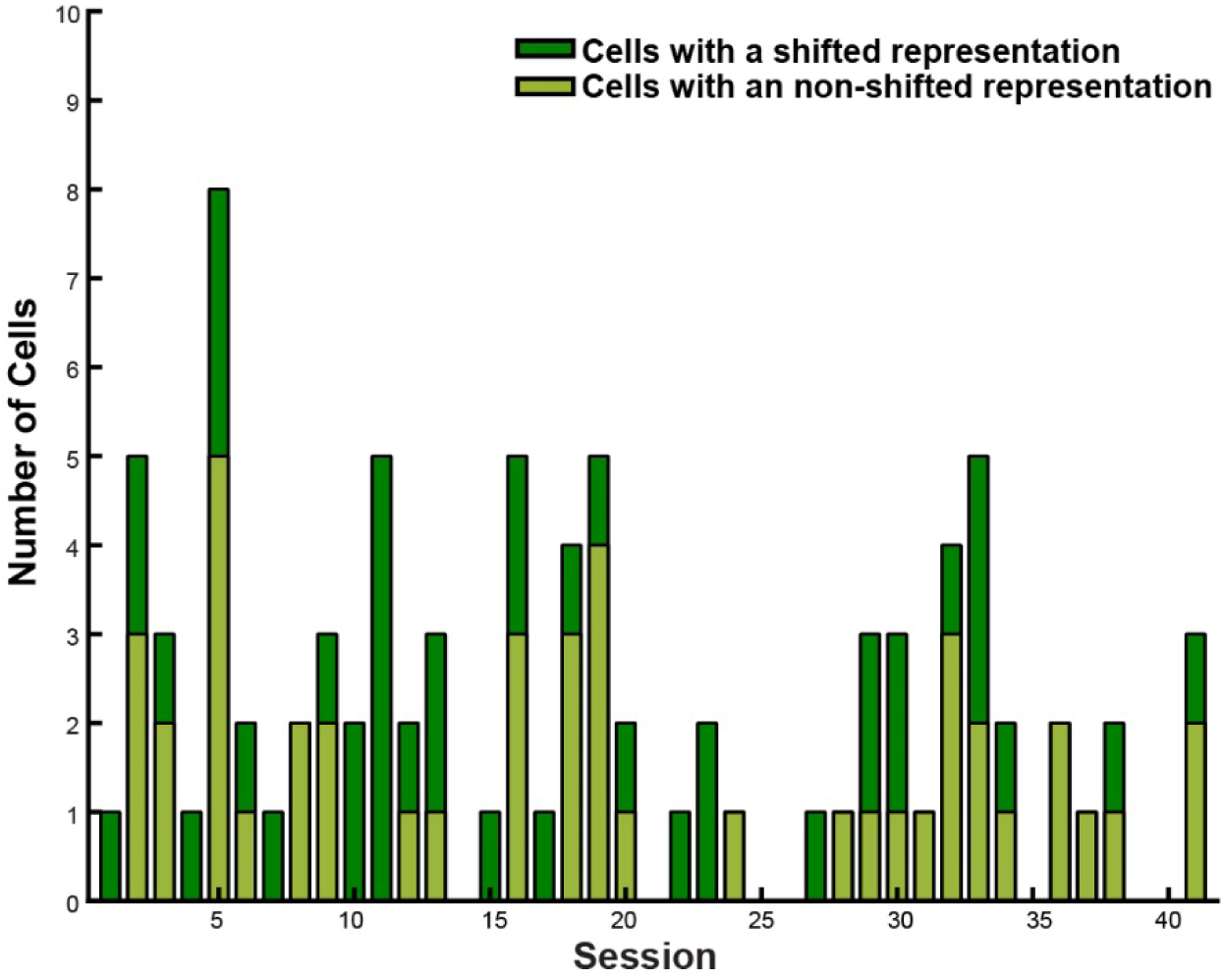
Neurons with different spatial reference frames were recorded simultaneously. The number of neurons that had a significantly consistent spatial representation across image window locations (n = 88) are shown for each recording session. Color indicates whether neurons exhibited a spatial representation that shifted with image window location.

In order to test whether a neural spatial activity shifted partially in the direction of the new image window location, spatial correlations for each neuron were computed for a range of partial spatial offsets (Figure 5). Each neuron’s rate maps from different image windows were correlated for eight spatial offsets that ranged between 0 and 100% of the image shift distance (4°). The incremental distance between different spatial offsets was one rate map spatial bin (0.5°). This analysis was performed three times using three different rate map smoothing methods (described above) in order to be agnostic about the “correct” smoothing. The resulting three correlation values for each offset were then summed in order to create one cumulative correlation vector per neuron across the range of spatial offsets, meaning that one cumulative correlation value corresponded to each offset. The offset with the highest value for each neuron is indicated in Figure 5 in red; the lowest value in blue. Variability was computed by re-sampling the given cell population with replacement for the same number of original cells to repeat the analysis 1000 separate times. Error bars in Figure 5 represent the middle 95% of values from these 1000 iterations.

**Figure 5.**
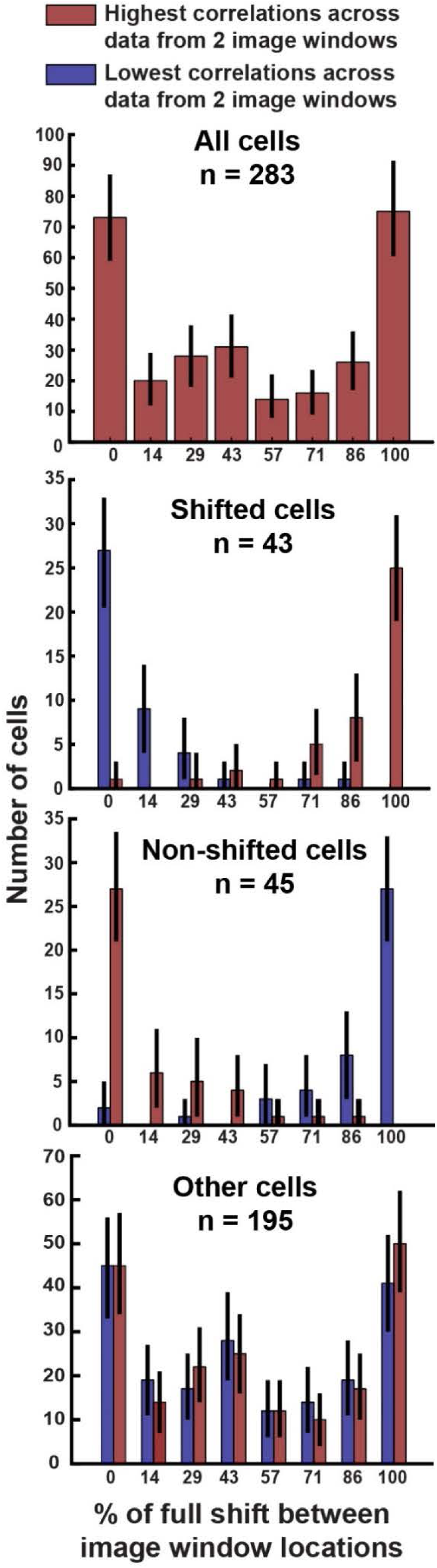
Neural activity is most spatially consistent for complete (instead of partial) shifts along with image window location. Cell rate maps from different image window location trial blocks were tested for partial shifting by computing the correlation values for a range of spatial offsets between 0 and 100% of the full shift of image window location. For each spatial offset, the red bars indicate the number of cells with their maximum spatial stability at that particular offset. This is shown for all cells (top row), cells that exhibited spatial activity which shifted with the image window location (2nd row, “Shifted cells”), cells that exhibited spatial activity which did not shift with the image window location (3rd row, “Non-shifted cells”), and cells which did not pass criterion to be categorized as Shifted or Non-shifted (bottom row, “Other cells”). Blue bars indicate the number of cells with their minimum spatial stability at each spatial offset. Error bars represent 95% confidence intervals of 1000 bootstrap iterations (re-sampling with replacement the cell population for the same number of original cells). Non-overlapping error bars confirm in each cell population that more cells have their maximum spatial correlation values at 0 and 100% shifts compared to partial shifts (p ≤ 0.05). See Materials and Methods for additional details on computing cell spatial stability.

### Saccade direction selectivity

Neurons meeting criterion for spatial stability were tested for saccade direction selectivity in order to avoid a potential confound within a rate map between gaze position firing fields and a saccade direction preference. If a neuron did show a saccade direction preference, then the neural data were tested to determine whether rate map stability was due to preferential firing for that saccade direction.

Saccade direction selectivity was tested in three different peri-saccade epochs: 100 ms leading up to the saccade, 100 ms centered on saccade onset, and a 100 ms period starting once the saccade completed. For each of these epochs, a neuron’s firing rate across different saccade directions was computed using a saccade direction bin 10 degrees wide, that incremented by 5 degrees. Using methods similar to (Killian et al., 2015), data were pseudo-randomly down-sampled so that each angular quadrant had the same number of trials. Any neurons with more than 10% of saccade direction bins lacking values were excluded from further analysis (1 neuron). If the down-sampled neuronal response showed significant (p ≤ 0.05) non-uniformity on the Rayleigh test and also no significant departure from a von Mises distribution on Kuiper’s test (p ≤ 0.05) (Berens and Valesco, 2009), the response was considered to be selective for saccade direction. No neurons showing significant non-uniformity on the Rayleigh test (0/16) showed a significant departure on the von Mises test. The preferred direction of a neuron was considered to be the direction with the maximum firing rate.

Rate map stability of neurons selective for saccade direction was re-computed after removing the data for the preferred saccade direction. Data were removed for saccades within 15 degrees of the preferred direction for any saccade epoch showing significant direction selectivity, and rate maps from these cut data were used to compute a new spatial correlation value. If this value was significantly lower than the original correlation (lower than or equal to the fifth percentile of 100 bootstrapped correlation values generated from the size-matched, down-sampled data from the original rate maps for p ≤ 0.05), than the spatial stability of the original rate maps was considered to be owed to the neuron’s saccade direction preference, and the compared rate maps were no longer deemed to show spatial stability.

### Image salience

Neurons with spatial stability were tested for a representational confound between gaze location and image salience by computing a spatial correlation (Spearman’s rank correlation coefficient) between the cell’s rate map and the image salience map of presented stimuli (Saliency Toolbox (Walther and Koch, 2006)). A correlation was considered significant if it was higher than the middle 95% of a distribution of bootstrapped correlation values between the image salience map and 1000 rate maps generated by shuffling the original rate map data.

### Predicting gaze location from neural activity

In order to assess the quality of the spatial information carried by the population as a whole, we used neural data from all 349 recorded neurons to predict gaze location. We first selected the two rate maps for each neuron that had the highest spatial stability as determined by the spatial correlation weighted by its percentile within shuffled correlations of the same data. Then, these two rate maps for each neuron were stacked with rate maps of other neurons in order to create two population rate maps (Figure 6A). In this way, a neuron could contribute either the first and second half of its data within one image window, or data from the 1st and 2nd image windows aligned to image window borders or to the screen borders. This process not only allowed an agnostic approach for targeting the most spatially consistent activity across all cells even if they were not categorized as spatial, but also permitted usage of cells with only data from one image window location. In order to be stacked, all rate maps were made to be the same size (the size of the screen area common to all image windows). The size of rate map spatial bins (0.5° × 0.5°) was not changed. The two population maps therefore reflected data from two different time periods for each neuron in the population.

**Figure 6.**
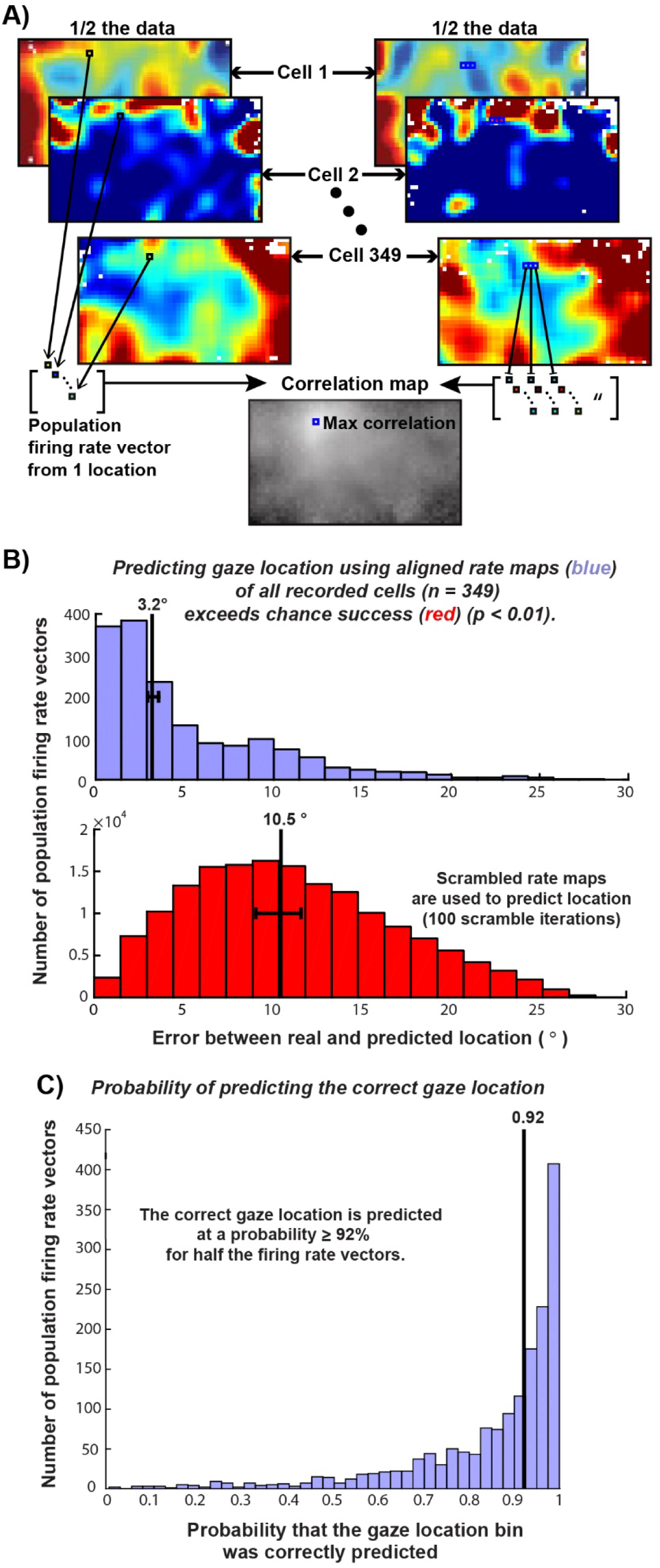
Gaze location can be predicted from population neural activity. (A) **Schematic of how gaze location was predicted for one population firing rate vector in a population firing rate map.** Each cell contributed data from two different time periods (two rate maps) in order to create two population rate maps, each with ∼half the data. Within the population map shown on the left, a small black square located in the same location on each cell’s rate map indicates the location of one gaze location spatial bin. The values of that gaze location bin location across rate maps constitute that spatial bin’s population firing rate vector, shown on the bottom left. The gaze position bin location of this vector is then predicted, by finding the highest correlation between the vector and all the vectors in the other population rate map containing the other half of the data, shown on the right. The bin location of the vector with the highest correlation was taken as the predicted gaze location. Across all predictions, each spatial bin (0.5° × 0.5°) was limited to being predicted once, meaning that if two different vectors from the map shown on the left both had their highest correlation with the same bin vector in the map shown on the right, then only one vector (with the higher correlation value) would be predicted to have that spatial bin location. (B) *Predicting gaze location of population firing rate vectors using aligned rate maps (blue) of all recorded cells (n = 349) exceeds chance success (red) (p ≤ 0.01).* Histograms show smaller (p ≤ 0.01) median error between the real and predicted gaze location when aligned neural activity was used to predict gaze location (top row) than when the prediction was made from scrambled rate maps (bottom row). Error was estimated from 100 iterations of predicting location of each firing rate vector from scrambled population rate maps. Top row: Vertical line indicates the median (3.2°), and the error bar indicates the entire range (100% confidence interval) of median error values generated by re-sampling the prediction error distribution with replacement 100 times. Bottom row: Vertical line indicates the prediction error expected in chance success (10.5°), marking the average median value of prediction error across 100 prediction iterations using scrambled population rate maps. The error bar indicates the entire range (100% confidence interval) of those median error values. (C) *Probability of predicting the correct gaze location using neural activity of all recorded cells (n = 349) is shown across all firing rate vectors.* Probability was determined by the rank of the correct gaze location among all potential locations ordered by their correlation value. In other words, if the correct gaze location bin had the highest correlation value across all potential gaze location bins, the probability of predicting it would be “1”, but if it were ranked 80^th^ out of 100 potential bin locations, then the probability of correctly predicting it would be 0.20).

The population firing rate vector of each spatial bin (0.5° × 0.5°) in one population map was then treated as a vector whose gaze location needed to be estimated (Figure 6A). This estimate was made by correlating the population firing rate vector in question with every spatial bin’s population firing rate vector in the other population rate map (Pearson’s correlation), and then choosing the bin with the highest correlation value. This chosen bin served as the estimate of gaze location for the vector in question. Because all rate maps of individual neurons were normalized between 0 and 1 before being added to the population rate map, general firing rate levels of individual neurons did not aid prediction. Each gaze location bin could be chosen only once, so a unique gaze location was predicted for every firing rate vector. If one gaze location was the best match for more than one vector, then that location was matched to the vector with which it was best correlated. Prediction error was computed as the Cartesian distance, in units of degrees of visual angle, between the actual location and predicted location of the population firing rate vector in question. The distribution of prediction errors for all 1623 firing rate vectors is shown in the top panel of Figure 6B. To estimate the variability of the median of this distribution, the distribution was re-sampled with replacement for the same number of predicted gaze locations to create a bootstrapped error distribution, and the 50^th^ percentile of this distribution was stored. This process was repeated 100 times to produce 100 bootstrapped values that represented the variability of the median prediction error, indicated by a horizontal error bar in the top panel of Figure 6B around the value 3.2°.

In order to compare the gaze location prediction to chance success, we predicted gaze location the same way as described above, except we scrambled the population rate maps by shifting each cell’s contributing rate maps (one “layer” within the population rate map) a random x-y distance. This resulted in population rate maps with the same general spatial structure as those used in the original analysis, except that the positions of firing fields were randomly shifted within each component cell’s rate map. Like above, the 50^th^ percentile of prediction error was used to describe the typical error of the prediction across vectors. This scrambling and subsequent prediction process was performed a total of 100 times in order to quantify the prediction error that would be expected if firing fields had no spatial stability. The 50^th^ percentile of prediction error for each iteration was stored, and these values represented the typical prediction error expected by chance. These 50^th^ percentile prediction error values were then compared between the scrambled and original data (Figure 6, 6-1 and 6-2). If the middle 95% of the error values from the original data were lower and did not overlap with the middle 95% of the error values from the scrambled data, then the original prediction was considered to have significantly lower error than would be expected by chance (p ≤ 0.05).

### Grid activity

Each cell was tested for grid activity by determining grid scores (Sargolini et al., 2006; Brandon et al., 2011; Killian et al., 2012) for each of its rate maps, the significance percentile of those grid scores, and the spatial stability across time of any map with a significant score. No rate map was considered to have significant grid activity if it lacked significant spatial stability in the relevant image space. For example, a neuron was considered to show shifted grid-like activity if it was one of the neurons categorized as having shifted spatial activity (described above) and it had a significant grid score relative to shuffled values for any image-aligned rate maps. Likewise, a neuron was considered to show non-shifted grid-like activity if it was one of the neurons categorized as having non-shifted spatial activity and it had a significant grid score in any screen-aligned rate maps. A neuron with data from only one image window was considered to show grid activity if it had a significant grid score and showed significant spatial stability across time across the halves of its data. A neuron was also considered to show grid activity if it had lacked stability across image windows, but had significant grid score for a spatially stable rate map in a single image window. All these tests were performed separately for each of the three different rate map smoothing methods described earlier.

The same tests were also performed to examine grid activity across smaller rate maps (Figure 7-1), except that those tests were intended to illustrate a conceptual point and were therefore less exhaustive. The aim was to test whether smaller rate maps, more comparable in size to past research from our laboratory (Killian et al., 2012), would yield a comparable proportion of neurons with grid activity. Each neuron’s rate maps were split across space along the two diagonals of the map (i.e., one rate map was divided into four rate maps with some overlapping data, see Figure 7-1), and then any map with grid-like activity (a grid score > 95^th^ percentile of 100 grid scores produced from that map’s shuffled data) that was also spatially stable was counted as a neuron showing stable grid activity for this smaller space. Neurons with stable grid activity from this analysis are reported separately with regard to this test, and not grouped in with the other analysis of grid activity that used the whole image-viewing area. Also, tests for grid activity across a smaller area differed in that they excluded rate maps smoothed adaptively, rate maps aligned to screen bounds from a single image window, and all cells with data from two image windows that only were significantly stable within, but not across, image windows.

Grid scores for each rate map were computed from its autocorrelation in two different ways, using the higher score as the final score (Killian et al., 2012). In the first method, the six closest peaks to the center peak of the autocorrelation were detected as the six bins closest to the center peak that each had a positive value higher than the surrounding twenty-four bins. The ringed area of the autocorrelation map that included these six peaks but excluded the central peak was then extracted, and the values for each radial location along the ring were averaged together to create one value for each position on a ring vector, and then this ring vector was correlated with itself at rotational offsets of 30°, 60°, 90°, 120° and 150°. The grid score was then calculated as the maximum correlation value of the 30°, 90° and 150° rotations (the angle rotations where a grid cell with 60-degree symmetry would be expected to have a low correlation) subtracted from the minimum correlation value of the 60° and 120° rotations (rotations where a cell with 60° symmetry would be expected to have a high correlation). The initial ring area extracted from the autocorrelation began at half the distance of the average peak distance from the center, and was only 2-bins wide. In subsequent iterations, the width of the extracted area grew by one bin, and the rotational values were repeatedly assessed to produce additional grid scores for fatter ring areas. The maximum grid score from all ring widths was taken as the grid score for this method. In the second method for computing grid scores that corrected for elliptical distortion of 60° symmetry (Brandon et al., 2011), the same process was repeated except the extracted ring area from the autocorrelation was adjusted to be an ellipse (the furthest peak was considered the major axis of the ellipse, or, in another full repetition of grid score calculation, the closest peak was considered the minor axis of the ellipse). The highest resulting grid score across the two methods was used as the final score for a rate map. A grid score of a given rate map was considered significant if it was greater than or equal to the 95^th^ percentile of synthetic grid scores produced from shuffling the data for that rate map 100 times (shuffling procedure described in *Spatial Stability* section).

## Results

### Consistent spatial activity of individual neurons

349 single entorhinal neurons (237 from Monkey WR and 112 from Monkey MP) were recorded across 41 sessions. Over the entire population, about 40% (137 / 349) of all recorded neurons exhibited a consistent spatial representation, either within a given location or across image locations. About one fourth of all neurons (77 / 349) consistently represented gaze position across trials in which the images were presented in the same location (top panels of Figures 2 and 3). Across trials in which images were presented in different locations, about one third of neurons with data from two image window locations (88 / 283) exhibited a consistent spatial representation. Half of these neurons (43 / 88) exhibited an allocentric spatial representation locked to the bounds of the image display window, signaling gaze location relative to the bounds of the image display. For example, the neuron shown in Figure 2 (also Extended Data Movie 1) shifted its spatial representation along with the shifted location of the image display window. The other half (45 / 88) of these neurons showed a spatial representation that did not shift with the location of the image window on the screen, suggesting that they were signaling gaze position in an egocentric or stationary reference frame (Figure 3 and Movie 2). These relative proportions were similar in each monkey (shifting spatial activity, Monkey WR: 53%, Monkey MP: 41%). Additional examples of neurons with consistent spatial activity are shown in Figure 2-1. Interestingly, neurons with different spatial reference frames, i.e., with shifting or non-shifting spatial activity, were often recorded simultaneously (Figure 4).

All neurons with spatial consistency were further analyzed to determine the extent to which any saccade-direction selectivity or salient image features may have contributed to observed spatial representations. 12% of spatial neurons (16 / 137) showed selectivity for a saccade direction, but all of these neurons maintained spatial stability (p ≤ 0.05) after removing the preferred saccade direction responses. Regarding salient image features, 1% of spatial neurons (2 / 137) showed a consistent central spatial firing that was correlated (p ≤ 0.05) with the central spatial bias of salient image regions.

In order to determine whether spatial activity shifted partially in the direction of a new image window location, spatial correlations for each neuron were computed for a range of spatial offsets between 0 and 100% of the 4° distance between image windows. Across all neurons, neural activity was most consistent across image window locations for complete (0 or 100%) rather than partial shifts (Figure 5, top panel). For the population of neurons with spatial representations that shifted along with image window location (“Shifted cells”), neural activity was most consistent when spatial stability was tested at a completely shifted offset (Figure 5, 2^nd^ row, red bars), and least spatially consistent at a non-shifted offset (Figure 5, 2^nd^ row, blue bars). The reverse was observed for the population of cells with spatial representations that did not shift along with the image window location (“Non-shifted cells”; Figure 5, 3^rd^ row). For example, the Non-shifted cell shown in Figure 3 exhibits spatial activity that is asymmetric, and therefore its spatial correlation for non-shifted, screen-aligned activity (r = 0.54) is much higher than the correlation when its activity is aligned to the image frame (r = 0.39).

While about one third of neurons recorded with two image locations (88 / 283) met criterion as spatially stable across different image window locations, it is clear from the bottom panel of Figure 5 that the remaining cells (n = 195) mimic the correlation pattern of the cells that passed criterion; their rate maps also have maximum correlation values at complete (0 or 100%) rather than partial shifts with image window location. Additionally, like the cells that passed criterion for spatial stability, about equal numbers of this cell population were maximally stable when perfectly aligned with the image window location or aligned with the screen. Including these neurons which exhibited maximum spatial stability at 0 or 100% of spatial offset, the majority of all recorded neurons (63%, 221 / 349) represented gaze location.

### Predicting gaze location from neural activity

In order to assess the quality of the spatial information carried by the population as a whole, we used neural data from all 349 recorded neurons to predict gaze location. A gaze location was predicted for each population firing rate vector in a population rate map (Figure 6A). When considering all 1643 population firing rate vectors in the population rate map, the size of the resulting prediction error was quite small; half of the vectors were predicted within 3.2° of their actual location (Figure 6B, top). The bottom panel in Figure 6B shows what prediction error would result by chance, with predictions based on scrambled population rate maps, in which each cell’s rate map was shifted circularly by a random × and y value (causing the firing fields to change location). The proportion of population firing rate vectors predicted within various amounts of error is shown in Figure 6-1. Even when only 10 neurons were used to predict gaze position, the median error was significantly lower than chance (Figure 6-2). When 40 neurons were used to make the prediction, the median prediction error was already ∼3° and as low as 2.5° for some groups of neurons tested (Figure 6-2).

### Grid activity

A proportion of neurons with stable spatial activity (13 / 137, 10%) demonstrated grid-like representations and had significant grid scores (Figure 7). These grid cells were, like the general population of spatially consistent neurons, about evenly split between those which had a spatial representation that shifted along with image window location (n = 6) and those that did not show a shift (n = 4). The remaining three grid cells either showed stable grid activity only across trials where the image window was in one location (n = 2), or could not be tested for spatial stability beyond one image window because data was only collected from one image window location (n = 1).

**Figure 7.**
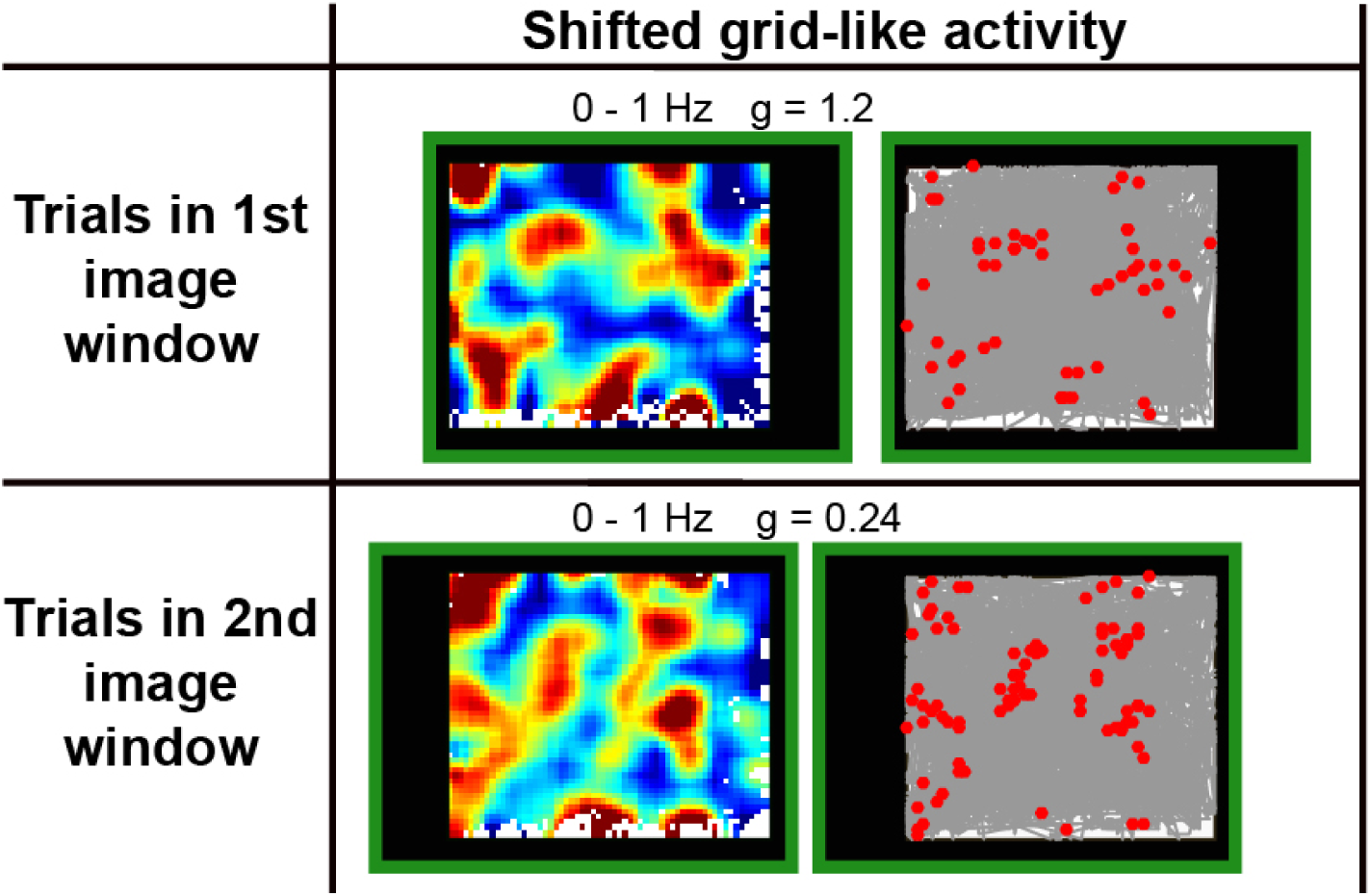
Grid-like activity can shift along with image window location. An example neuron is shown that yielded a significant grid score (p = 0.04) for a spatially stable rate map across two image window locations. The neuron’s spatial activity shifted with the location of the image window on the screen (r = 0.4 spatial correlation between rate maps from two image window locations aligned to image window bounds, p = 0.01). A movie of spikes occurring for this neuron as gaze position moves over the screen can be found in Extended Data (Movie 3). Plotting schematic is the same as Figure 2. The grid score for each rate map (g) is indicated on the top of the map. Rate maps of the shifted activity are the same size as presented images (30° × 25°).

Notably, grid cells accounted for a small proportion of the total recorded neurons (13 / 349, 4%, without pre-selecting spatially consistent neurons) and contrasts with the larger proportion (12%) identified in a previous study from our laboratory (Killian et al., 2012). Although only about one fourth of our recorded neurons (n = 95 / 349) were located in the superficial layers where grid cells are predominantly found in rodents (Figure 4-1), we suspect that this is not primarily responsible for our low yield of grid-like activity, since the previous study identified grid cells in both superficial and deep layers. Rather, we speculate that our observation of a lower proportion of neurons with grid activity is attributable to the fact that we measured spatial activity across a much larger viewing area than in the previous study, with a viewing area subtending 4 to 7 times more area in degrees of visual angle. Although a smaller area certainly provides a lower bar for detecting grid activity simply because a grid pattern need only be consistent across a smaller area, grid patterns have been shown to lose consistency across very large spaces in rodents (Stensola et al., 2015).

To examine whether a smaller viewing area would result in more reliable grid activity, we assessed grid activity and its stability for each neuron for half the area of its rate maps by dividing each rate map along each diagonal (Figure 7-1, Movies 4 and 5). Consistent with qualitative inspection, this analysis revealed a larger proportion of neurons with significantly stable grid activity (47 / 349 cells, 13%), which was comparable to the proportion reported in earlier work (Killian et al., 2012). Importantly, even within the smaller area, neurons with stable grid activity had more than six firing fields (Figure 7-1).

## Discussion

Entorhinal neurons including grid cells, border cells, and head direction cells represent body position relative to world features as rodents actively explore their environment with locomotive movement. In monkeys, similar spatial representations have been identified during visual exploration with eye movement; however, the frame of reference for these spatial representations in primates was unknown. In particular, it was unclear whether entorhinal neurons code gaze position relative to visual world features, similar to the allocentric activity identified in rodents. Also, the proportion of neurons with spatial representations had never been examined. Accordingly, this work sought to assess the extent of primate entorhinal neurons with spatial representations, and identify the spatial reference frame used, by recording spiking activity while monkeys freely viewed images displayed at different locations.

Our results revealed that a majority of primate entorhinal neurons represent gaze position. Roughly half of these neurons fired consistently when a monkey fixated specific locations within an image display window, even when the location of the image window was moved within the screen, demonstrating that individual primate entorhinal neurons can reflect an allocentric, visual frame of reference. Because not all simultaneously recorded neurons exhibited spatial firing that moved along with the location of the image display window, these results demonstrate that co-active neurons do not necessarily code gaze position within the same spatial frame of reference. Importantly, while the firing fields we observed most often exhibited an irregular layout across space, the spatial activity across the neural population was stable and specific enough to allow for successful prediction of gaze position, with the low median error of 2.5° between actual and predicted location.

### Co-active neurons with distinct reference frames

We found that the reference frame can differ across simultaneously recorded cells. These findings are consistent with some reports of rodent entorhinal cells non-coherently altering their spatial responses to environmental change (Savelli et al., 2008; Stensola et al., 2012), but stand in contrast to other reports of coherent spatial responses across cells to environmental change (Hafting et al., 2005; Fyhn et al., 2007; Solstad et al., 2008; Savelli et al., 2017). One explanation for the discrepancy across studies could be the variability in the strength of the environmental change (Jeffery, 2011). Cells might respond to environmental change independently of one another only when the change is subtle, which is perhaps a condition met in our experiment. Supportive of this idea, changing only the odor of an environment resulted in partial remapping in the rodent hippocampus, where only some cells within a population of simultaneously recorded cells changed their spatial activity (Anderson and Jeffery, 2003). Likewise, changing the location of a key landmark in a foraging task rich with other landmarks revealed that co-active hippocampal cells could show spatial firing fields relative to that landmark or the unchanging landmarks of the room (Gothard et al., 1996). Such mixed-reference frame hippocampal activity is potentially a downstream reflection of entorhinal cell activity. In line with this idea, inactivation of the medial entorhinal cortex in rodents can cause partial remapping in hippocampal CA1 neurons (Rueckemann et al., 2016), indicating that spatial responses of hippocampal cells can be directed by entorhinal input. Taken together, these results suggests that both entorhinal and hippocampal neurons can sustain multiple reference frames at one time across the population, and may do so in concert as connected sub-networks across brain areas.

An important caveat is that we did not exhaustively test spatial responses relative to all possible shifts in reference frame. Specifically, we did not chronically monitor cell activity over multiple experimental sessions to determine whether individual cells switch reference frames over time. In addition, we did not examine the neuronal responses to shifts in other possible perceptual reference frames, such as the screen itself or the room. Earlier work in the primate hippocampus found only a minority of neurons coded for gaze position relative to the body axis in true egocentric coordinates (Feigenbaum and Rolls, 1991).

### Prevalence of irregular spatial activity

We designed this study to enhance our ability to identify grid cell activity by using larger visual images than our previous study, thus allowing for more firing fields within each image (Killian et al., 2012). However, we instead observed a dominance of irregular spatial cells that informs our understanding in several ways. First, the small amount of cells with coherent grid activity over the large image space could be viewed as consistent with results in rodent in which grid representations were influenced by local features, thus especially distorting the coherence of the grid pattern across large enclosures (Stensola et al., 2015). Congruent with this idea, the data from the present study demonstrate that some irregular spatial cells have significant grid-like activity within a restricted region of the visual display (Figure 7-1). Secondly, the dominance of irregular spatial cells suggests that they play an important role within the circuit. In rodents, about 10% of entorhinal cells in the superficial layers of medial entorhinal cortex are grid cells (Tang et al., 2014; Sun et al., 2015), while irregular spatial cells are widespread, comprising as much as 50 - 70% of superficial layer medial entorhinal cells (Sun et al., 2015; Diehl et al., 2017), and they likely constitute a major input to the hippocampus (Zhang et al., 2013). Irregular spatial cells also show persistent spatial responses, along with place cells in the hippocampus, even when theta oscillations and grid cell patterns are diminished by septal inactivation (Brandon et al., 2011; Koenig et al., 2011), indicating that irregular cells may play an important role in sustaining the spatially specific firing of place cells in the hippocampus (Poucet et al., 2013). The theoretical utility of cells with irregular fields for self-localization in large environments has been recently highlighted (Hedrick and Zhang, 2016). Here, our neural data, dominated by irregular spatial activity, was used to infer gaze position with a high degree of spatial resolution (within 2.5° of actual gaze position). Interestingly, this spatial resolution resonates ecologically; it is about the limit of visual space within which visual detail can be extracted during a single fixation (Findlay and Gilchrist, 2003).

### Entorhinal gaze position signal is not a traditional visual nor motor response

We identified a large portion of entorhinal neurons that represent gaze position. However, it is important to note that this entorhinal representation of gaze position is distinct from the traditional eye-centered responses observed in early visual cortex and oculomotor areas. Specifically, while a neuron in a visual or oculomotor area responds to a confined region of eye-centered space within the contralateral visual field (its response field), we observed that entorhinal neurons selectively fired when the monkey fixated multiple, locations within a 30°-wide image window, with highly variable scan paths across over one hundred images in each session. In addition, entorhinal neurons showed a lack of sensitivity to image content, firing consistently for fixation locations occupied by a large range of perceptually distinct images. These entorhinal properties are perhaps unsurprising given that the entorhinal cortex receives extensive input from the perirhinal cortex (Van Hoesen and Pandya, 1975; Suzuki and Amaral, 1994), where cells have large, bilateral receptive fields (Desimone and Gross, 1979).

Another notable distinction in the current research is the use of a free-viewing paradigm instead of the fixation-based paradigms traditionally used to measure visual and eye-movement responses in primates. The free-viewing paradigm used here is in many ways analogous to the free-foraging paradigms used to assess entorhinal spatial responses in rodents, and allows for cross-species comparisons. Moreover, this naturalistic, exploratory free-viewing paradigm was instrumental in identifying spatial representations in the primate entorhinal cortex (Killian et al., 2012; Killian et al., 2015), and this kind of paradigm has been shown to be sensitive to damage to medial temporal lobe structures in primates (Pascalis and Bachevalier, 1999; Ryan et al., 2000; Zola et al., 2000; Smith and Squire, 2008; Hannula et al., 2010).

### Conclusion

The present results provide evidence that entorhinal cells can code gaze position in a visual reference frame that is spatially broad and insensitive to image content. Such large, visual reference frames could be used to produce eye movements from memory that use a remembered environment as a frame of reference. Specifically, some eye movements may be guided by gaze position relative to the remembered structure of an environment, such as in natural behavior when people shift gaze to a target outside the current field of view (Land et al., 1999; Hayhoe et al., 2003). Recent work in patients with medial temporal lobe damage suggests a strong role for this brain region in the rapidly acquired memory of the spatial layout in a visually presented scene (Urgolites et al., 2017). How the entorhinal cortex, as part of a memory system, uses these spatial signals to produce adaptive behavior is a fascinating question, and future studies are necessary to advance our understanding of the neural mechanisms by which memory guides viewing behavior (Meister and Buffalo, 2016).

## Author Contributions

M.L.R.M. and E.A.B. designed the experiment, peformed the surgeries, and wrote the paper. M.L.R.M. collected and analyzed the data.

## Acknowledgments

This work was supported by NIH 2R01MH080007, R01MH093807 and NIMH P51 OD010425 awarded to E.A.B. We are grateful to Laura Kakalios, Kiril Staikov, Megan Jutras and Kelly Morrisroe for assistance with animal training and handling, to Nathan Killian for experimental advice and supplying example Matlab code for analysis, and to Jon Rueckemann for helpful discussion regarding the analysis of gaze position prediction. The authors declare no competing financial interests.

## Extended Data

### Movie 1. Related to Figure 2

The movie shows spiking activity from the neuron shown in Figure 2 as eye position moves over the screen. Data are shown separately for trials in which the monkey viewed images presented at two separate screen locations. Activity is aligned to image bounds.

### Movie 2. Related to Figure 3

The movie shows spiking activity from the neuron shown in Figure 3 as eye position moves over the images. Data are shown separately for trials in which the monkey viewed images presented at two separate screen locations. Activity is aligned to screen bounds.

### Movie 3. Related to Figure 7

The movie shows spiking activity from the neuron shown in Figure 3 as eye position moves over the screen. Data are shown separately for trials in which the monkey viewed images presented at two separate screen locations. Grid-like activity is aligned to image bounds.

### Movie 4. Related to Figure 7 and Figure 7-1

The movie shows spiking activity from the neuron shown in Figure 7-1 panel A as eye position moves over the screen. Data were collected from only one image window location. First and second half of trials are shown separately. Grid-like activity is shown over half of the image space.

### Movie 5. Related to Figure 7 and Figure 7-1

The movie shows spiking activity from the neuron shown in Figure 7-1 panel B as eye position moves over the screen. Data are shown separately for trials in which the monkey viewed images presented at two separate screen locations. Grid-like activity over half of the image space is aligned to image bounds.

**Figure 1-1.**
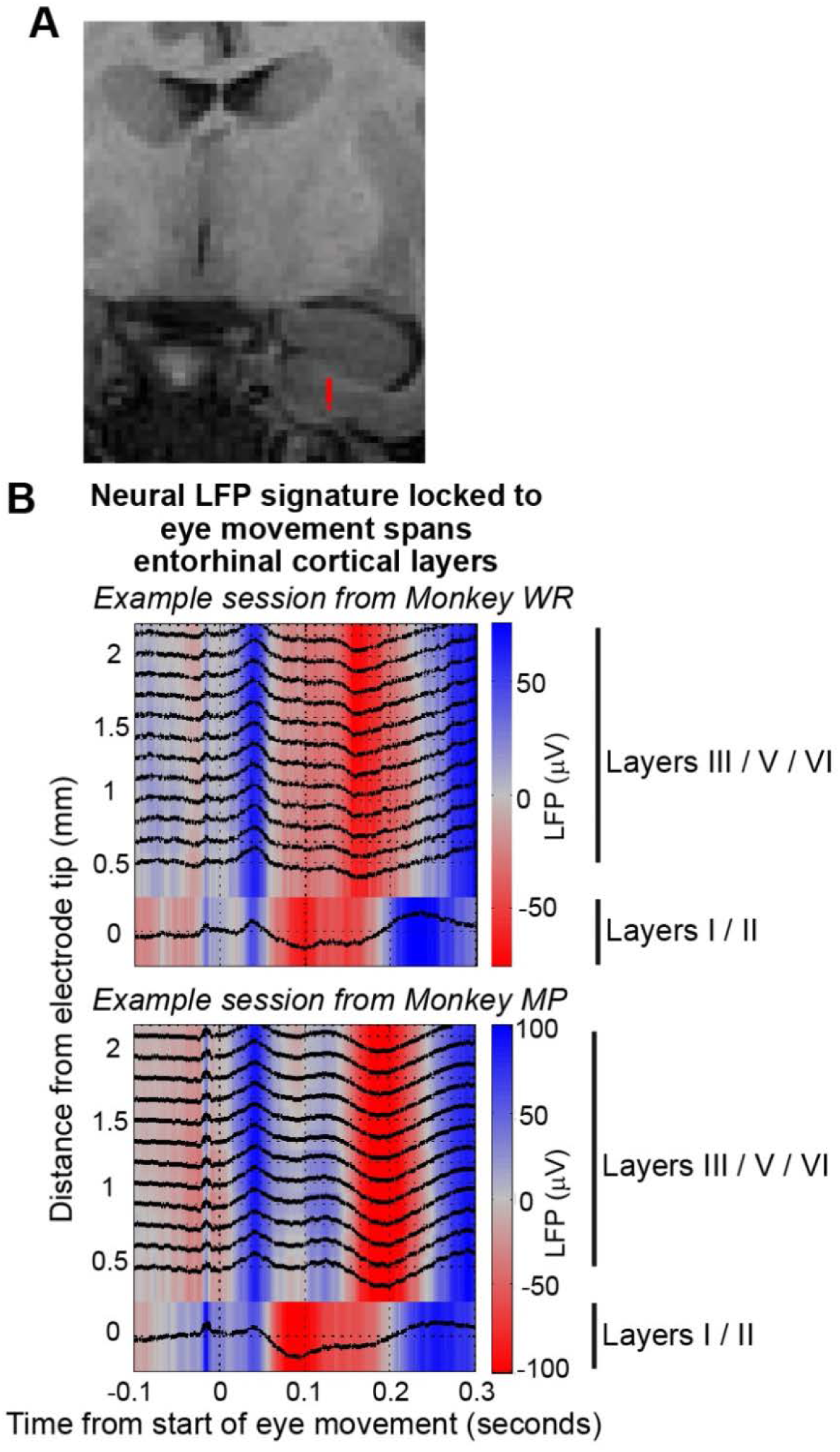
Recording location within the entorhinal cortex. **(A)** Estimated position of recording channels in the entorhinal cortex in one recording session is shown in red on a coronal MRI. **(B)** Successful targeting of the entorhinal cortex was further confirmed for each session by the electrophysiological signature of the local field potential (LFP) across cortical layers locked to eye movement (Killian et al., 2012; Killian et al., 2015). An example of this signature is shown from a session in each monkey. Black lines are individual average LFPs for each of the 13 recording channels. The amplitude has been normalized to the maximum over all LFPs. Data are aligned to onset of eye movement.

**Figure 2-1.**
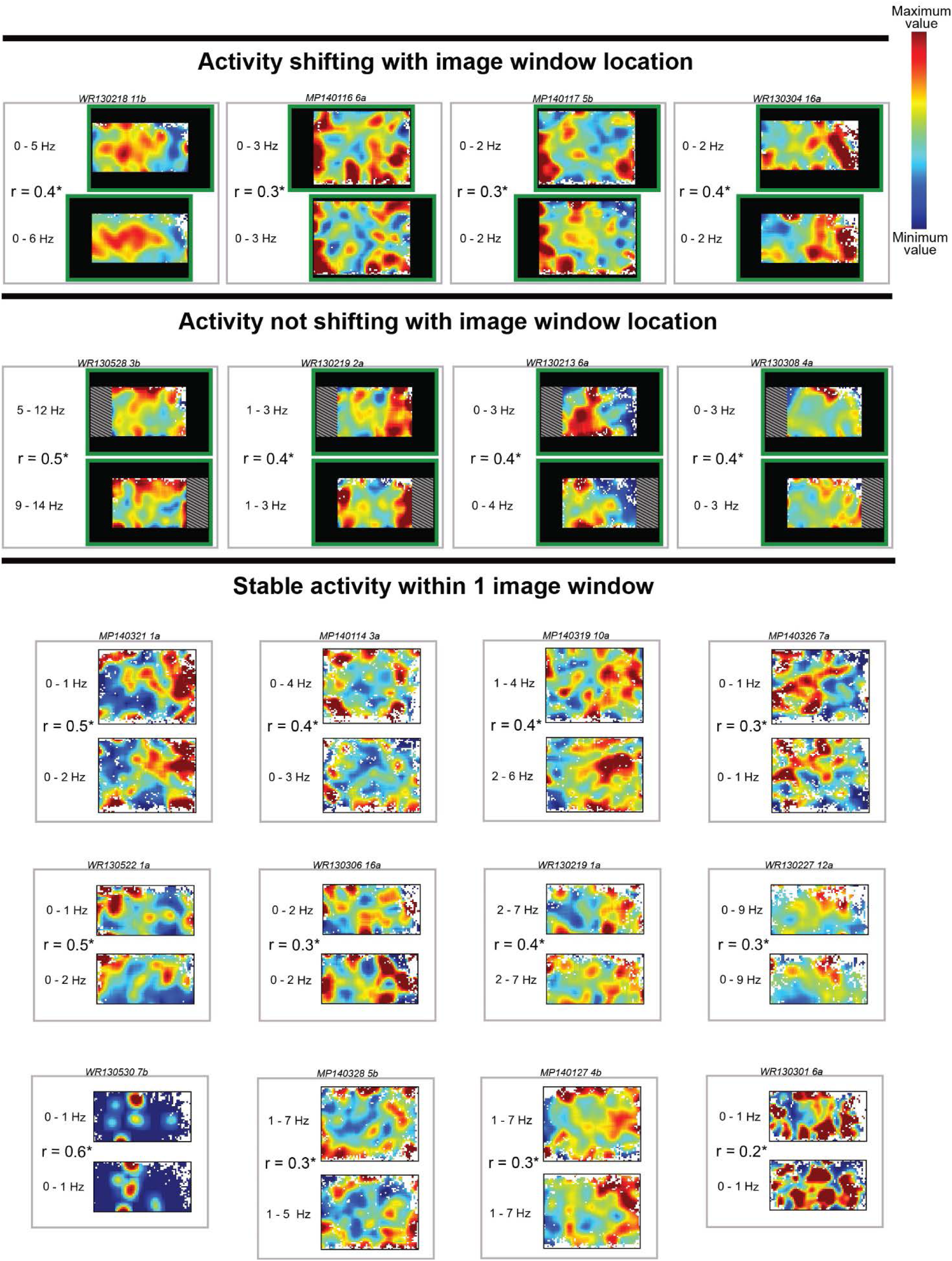
Additional example neurons showing stable spatial activity. Rate maps of single neurons (n = 20) with stable spatial activity are shown. Each gray rectangle shows the activity of one cell, within which the top and bottom rate map respectively show data from different portions of the recording session. For the top 8 neurons, the data are split between the different image window locations, and for the bottom 12 neurons, the data are split between the 1^st^ and 2^nd^ half of viewing within a single image window location. Warm colors indicate higher firing rate values; the firing rate scale reported to the left of each rate map corresponds to the maximum and minimum of the color bar (top right). Green rectangles indicate screen bounds. Crosshatching indicates the part of image window rate map that is not shown for visual clarity of non-shifting activity. The spatial correlation between each pair of rate maps is indicated by “r”, and followed by a “*” to indicate a significance of p ≤ 0.05. The top section shows cells that shifted spatial activity with image window location (rate maps are aligned to the image bounds). The middle section shows cells that did not shift spatial activity with image window location (rate maps over common image space are aligned to screen bounds). The bottom section shows cells with spatial activity that was stable within a single image window (rate maps aligned to image window bounds).

**Figure 4-1.**
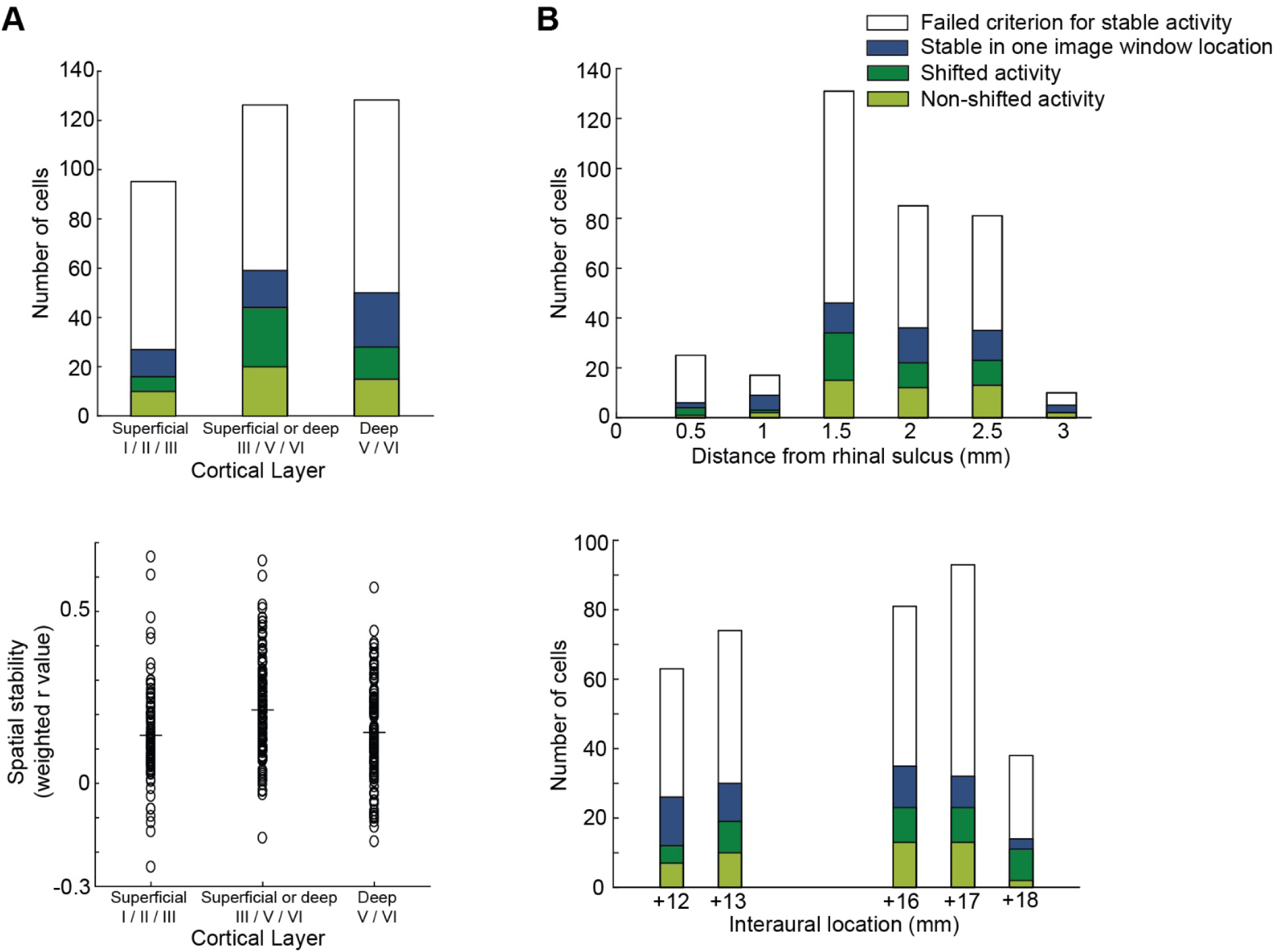
Anatomical distribution of spatial activity within the entorhinal cortex. Spatial activity did not appear to vary systematically across the EC. The proportion of cells with spatial activity did not vary distinctly across cortical layer (A), distance from the rhinal sulcus (B, top), or distance along the anterior-posterior axis (B, bottom). Colored portions of bar graphs indicate the proportion of cells that exhibited different types of spatial stability or spatial reference frame (see legend at top right). In the bottom panel of A, spatial stability for each cell is shown as a function of cortical layer. Spatial stability for a cell that did not pass criterion as a spatial cell is computed as the highest spatial correlation value for the cell across its rate maps (within or across image window locations) multiplied by its significance percentile (between 0 and 1). For cells with significant spatial stability (cells whose significance percentile is > 0.95), the highest significant correlation value was used. Black horizontal lines indicate the mean value of each cortical layer group.

**Figure 6-1.**
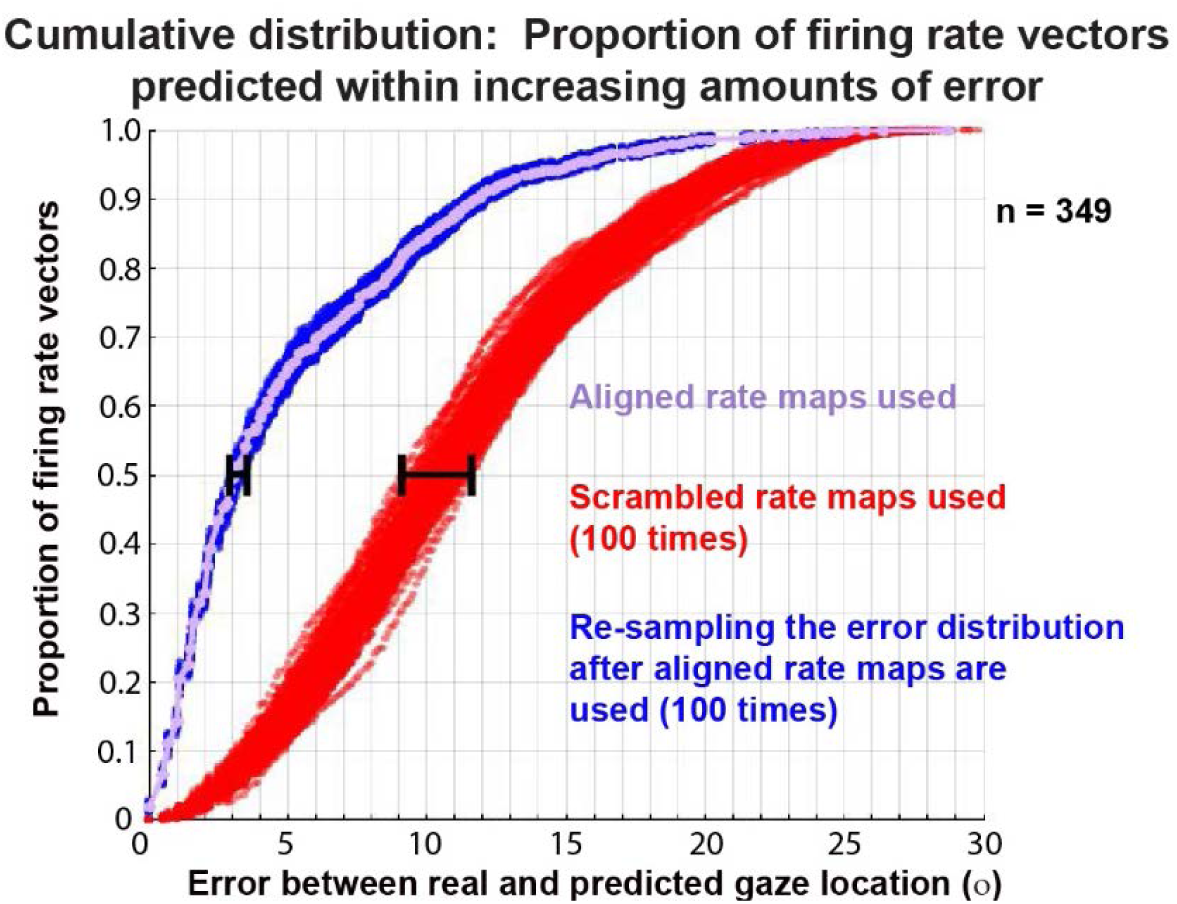
Predicting gaze location using population rate maps of all recorded neurons (n = 349) This cumulative distribution shows the proportion of population firing rate vectors as a function of the amount of error between real and predicted location of gaze, and demonstrates that gaze location is predicted more accurately when aligned neural activity is used to make a prediction (violet) than when scrambled rate maps are used (red, 100 scrambling iterations). Blue shows variability of the violet prediction error distribution, and represents 100 cumulative distributions from resampling the original error values 100 times with replacement. Horizontal error bars are identical to those shown in the histograms in Figure 6, and indicate the variability (100% confidence interval) of the 50^th^ percentile of prediction error.

**Figure 6-2.**
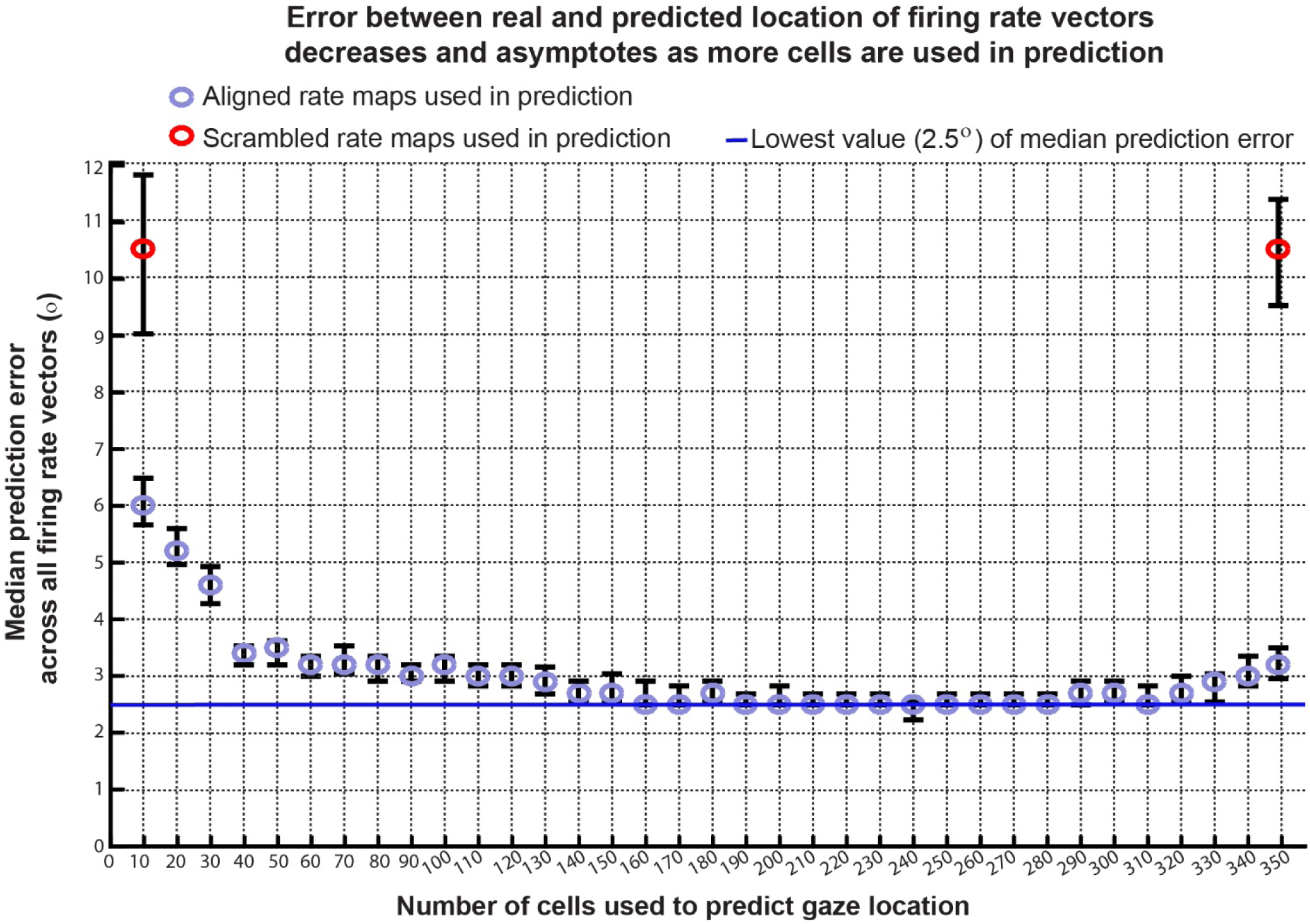
The activity of entorhinal neurons can be used to predict gaze location. The error between real and predicted location of population firing rate vectors (violet) decreases and asymptotes as more cells are used in the prediction. For comparison, the error of prediction using scrambled population rate maps is also shown (red). Cells were added in order of their spatial correlation weighted by significance percentile, so that the cells with the highest values were used first. The blue line indicates the lowest value (2.5°) of median prediction error. The asymptotic level of prediction error at ∼2.5° approaches the resolution of our eye position measurements, which is 0.5° at best (the width of our rate map spatial bins) across all sessions and visual space. Error bars indicate a 95% confidence interval from bootstrapped values. Bootstrapped error values for the aligned neural rate maps arise from resampling (with replacement) the original error distribution across all firing rate vectors for the original number of firing rate vector predictions, generating a 50^th^ percentile error from that resampled distribution, and then repeating this process 1000 times. Bootstrapped error values for the scrambled neural rate maps arise from taking the 50^th^ percentile of the error distribution after all firing rate vectors have been predicted using a pair of scrambled population rate maps, and then repeating this process 100 times. This latter process is computationally time-consuming, but the highly similar values of prediction error for scrambled rate maps for extreme values of the abscissa (when 10 or 349 cells are used in the prediction) suggests that ∼10.5° is the asymptotic error value for random prediction, and the rest of the “unfilled” intermediate portion between the two red points is the likely same value. Spatial bins were excluded from the prediction analysis if they had data from 5 neurons or less, which meant that at most 1% of the bins were excluded in predictions using 30 neurons or less.

**Figure 7-1.**
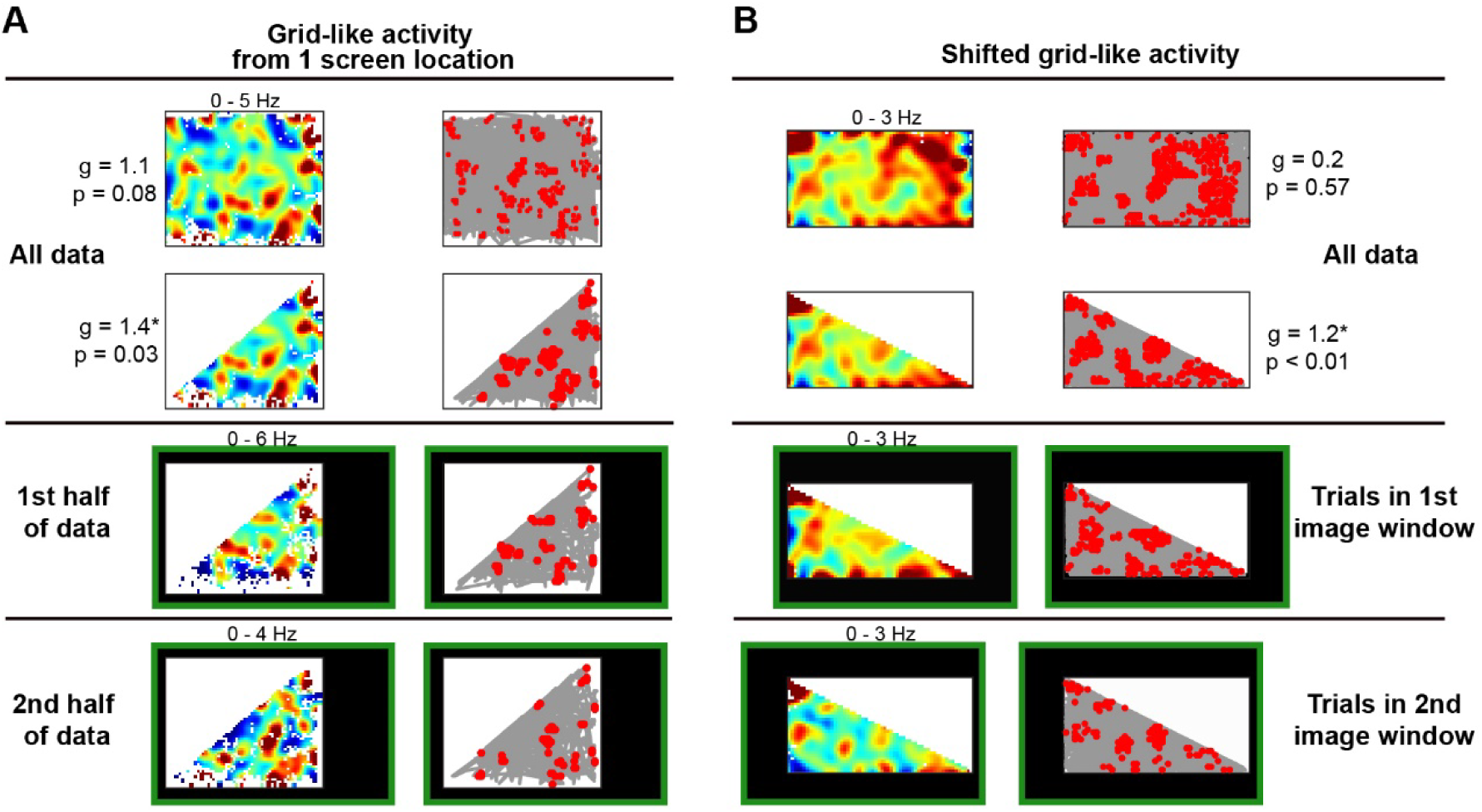
Neurons with non-significant grid scores can show stable, significant grid activity in part of their rate maps. Two example neurons shown (one cell in A, one cell in B) failed to reach significance for grid activity with their full rate maps (top row), but did show significant, stable grid activity when a smaller portion (half) of the rate map was used (2^nd^ row). g: grid score; asterisk indicates that the score is significantly higher (p ≤ 0.05) than bootstrapped grid scores from the same rate map when the spike train was synthetically displaced along the eye trace. Plotting schematic for each rate map and corresponding eye trace is identical to Figure 2. Movies of spikes occurring for these example neurons as eye position moves over the screen can be found in Extended Data Movies 4 and 5 for the neuron shown in A and B respectively. (A) Data were collected from only one image window location. The spatial activity of half the rate map was stable between rate maps (r = 0.2, p = 0.03) of the 1^st^ and 2^nd^ halves of the data (bottom two rows). Size of rate map in top row is 30° × 25°. (B) Cell spatial activity shifted with the location of the image window (r = 0.4, p ≤ 0.01 between rate maps from the 1^st^ and 2^nd^ image window location shown in bottom two rows). Rate map in top row is 30° × 15°.

